# Morphine exposure bidirectionally alters c-Fos expression in a sex-, age-, and brain region-specific manner during adolescence

**DOI:** 10.1101/2021.04.29.441958

**Authors:** C Figueroa, H Yang, J DiSpirito, JR Bourgeois, G Kalyanasundaram, I Doshi, SD Bilbo, AM Kopec

## Abstract

Drug and alcohol use during adolescence is common, and data in both humans and preclinical animal models clearly indicate drug exposure during adolescence increases the risk of substance use and other mental health disorders later in life. Adolescence is a period of social, emotional, and cognitive development, and is characterized by increased exploratory behavior, risk-taking, and peer-centered social interactions. These are thought to be behavioral manifestations of developmental plasticity in ‘reward’ regions of the brain. Human data indicate that adolescence is not a unitary developmental period, but rather different neural and behavioral sequelae can be observed in early vs. late adolescence. However, most studies with rodent models examine a single adolescent age compared to a mature adult age, and often only in males. Herein, we sought to determine whether the acute response to the opioid morphine would also differ across adolescence, and by sex. By quantifying c-Fos positive cells, a proxy for neural activity, at different stages during adolescence (pre-, early, mid-, and late adolescence) and in multiple reward regions (prefrontal cortex, nucleus accumbens, caudate/putamen), we determined that acute morphine can either reduce or increase c-Fos expression dependent on adolescent age, sex, and brain region. These data suggest that heterogeneity in the consequences of adolescent opioid exposure may be due to the interaction between age- and sex-specific developmental profiles of reward processing in individual brain regions. In future studies, it will be important to add age within adolescence as an independent variable to fully capture the consequences of healthy or abnormal reward-related neural development.

## Introduction

Drug and alcohol use during adolescence is common [1], and data in both humans and preclinical animal models clearly indicate drug exposure during adolescence increases the risk of substance use and other mental health disorders later in life [2–4]. In fact, even short-term prescription opioid use after a surgery or injury during adolescence can increase opioid misuse later in life [5–7], which may contribute to the current opioid epidemic. Adolescence is a period of social, emotional, and cognitive development, and is characterized by increased exploratory behavior, risk-taking, and peer-centered social interactions; these are thought to be behavioral manifestations of developmental plasticity in ‘reward’ regions of the brain, including the prefrontal cortex (PFC) and ventral and dorsal striatum (nucleus accumbens (NAc) and caudate/putamen (CPu), respectively) [8–12]. Human and rodent data indicate that adolescence is not a unitary developmental period, but rather different neural and behavioral sequelae can be observed in early vs. late adolescence, which can further be sex-specific [8, 13, 14]. The interaction between adolescent drug use and sex-specific neural development may explain some of the sex-specific presentation of substance use and other mental health disorders later in life [15, 16]. However, most studies with rodent models examine a single adolescent age compared to a mature adult age, and often only in males. We recently determined that there are unique, sex-specific molecular and behavioral profiles within adolescence in rats at postnatal days (P)22 (pre-adolescence), P30 (early adolescence), P38 (mid-adolescence), and P54 (late adolescence) [14]. Building on these data, herein we sought to determine whether the acute response to the opioid morphine would also differ across adolescence, and by sex. By quantifying c-Fos positive cells, a proxy for neural activity, in the PFC, NAc, and CPu after saline or morphine injections, we determined that acute morphine can both increase and reduce c-Fos expression dependent on adolescent age, sex, and brain region. These data suggest that heterogeneity in the consequences of adolescent opioid exposure may be due to the interaction between age- and sex-specific developmental profiles of reward processing in individual brain regions.

## Methods

### Animal model

Adult male and female Sprague-Dawley rats were purchased to be breeding pairs (Harlan/Envigo), and were group-housed with ad libitum access to food and water. Colonies were maintained at 23°C on a 12:12 light:dark cycle (lights on at 07:00) and cages were changed once per week. Females were separated from males at 20 days post-pairing. Litters were culled to a maximum of 12 pups on P2-5, and at P21 pups were weaned into same sex pair-housing. At the ages indicated (P22, P30, P38, or P54), rats were subcutaneously injected with either 3mg/kg free base morphine sulfate (NIDA Drug Supply) or equivalent volume of sterile saline. Rats were euthanized by CO_2_ anesthesia and exsanguination 90mins after injection, when c-Fos induction is maximal [17–19]. Experiments and animal care were approved by the Institutional Animal Care and Use Committees at Massachusetts General Hospital and Albany Medical College.

### Tissue collection

Tissues were transcardially perfused with 0.9% saline followed by 4% paraformaldehyde, and then brains extracted and post-fixed for an additional 24 hrs at 4°C. Brains were then cryoprotected in 30% sucrose in 0.1M PB with 0.1% sodium azide until they were saturated, and then frozen in molds filled with O.C.T. on a slurry of isopentane and dry ice. Brains were cryosectioned at 30μm (1:8 series) and stored in 0.1M PB with 0.1% sodium azide until immunohistochemical processing.

### Immunohistochemistry (IHC)

A detailed IHC procedure prior to antibody incubations can be found in Kopec et al. 2018 [14]. Sections were incubated in mouse anti-c-Fos (Novus Biologicals NBP2-50037; 1:5000) overnight at 4°C, rinsed, and then incubated in donkey anti-mouse biotin secondary antibodies (Jackson Immuno #715-065-150; 1:500) for 2 hrs at room temperature. Tissue was rinsed, incubated with Vectastain ABC kit, rinsed, and then resolved with ImmPACT DAB Substrate Kit (both from Vector Laboratories) per manufacturer’s instructions. Sections were mounted onto gelatin-subbed slides, baked at 50°C for 2 hrs, and then dehydrated and defatted with ethanol and Histochoice Clearing Agent (Sigma-Aldrich), respectively. Slides were coverslipped with DPX mountant (Sigma-Aldrich) and imaged at 20X with a Hamamatsu Nano Zoomer 2.0-RS Slide Scanner.

### c-Fos quantification

The number of c-Fos positive nuclei within each region of interest (ROI) were quantified with our semi-automated analysis pipeline, described in detail in Bourgeois et al. 2020 [20]. Briefly, each hemisphere is aligned to the closest figure in Paxinos & Watson’s Rat Brain Atlas [21]. Landmarks are placed on the brain image to indicate corresponding areas on the atlas, including superficially around the brain perimeter and deep white matter structures, e.g. the corpus callosum and anterior commissure. These landmarks are used to alter the brain image to align more accurately with the atlas. Pre-coded ROIs from the brain atlas are then automatically overlaid on the image, and c-Fos positive nuclei are counted. Image alterations do not change the pixel size of c-Fos positive nuclei, and permit comparison of data across different brain sizes (i.e. developmental changes and sex differences) [20]. The number of c-Fos positive nuclei is normalized to the area of the ROI, as ROI areas vary according to Bregma. Figs. 8-13 (Bregma +4.2 - +2.52) were analyzed for PFC, Figs. 10-26 (Bregma +3.24 - +0.84) for NAc, and Figs. 12-26 (Bregma +2.76 - +0.84) for CPu.

### Statistics

For each ROI, c-Fos positive cells per ROI area values across Bregmas were treated as individual observations so as not to obscure innate biological variability. Data points greater than two standard deviations above or below group means were excluded as outliers. Because there were differences in c-Fos expression across adolescence in saline-treated rats, for the main figures (Fig. 1-6), we normalized morphine data to the average saline group data. Three-way ANOVAs (sex X age X drug) were conducted for each ROI, and when significant (*p*<0.05), Šídák’s multiple comparisons tests were performed post-hoc. The results of the post-hoc tests are indicated by asterisks in Figs. 1-6. Data were analyzed, and histograms created, with GraphPad Prism 9. Data are depicted as average ± standard error of the mean.

**Fig. 1:**
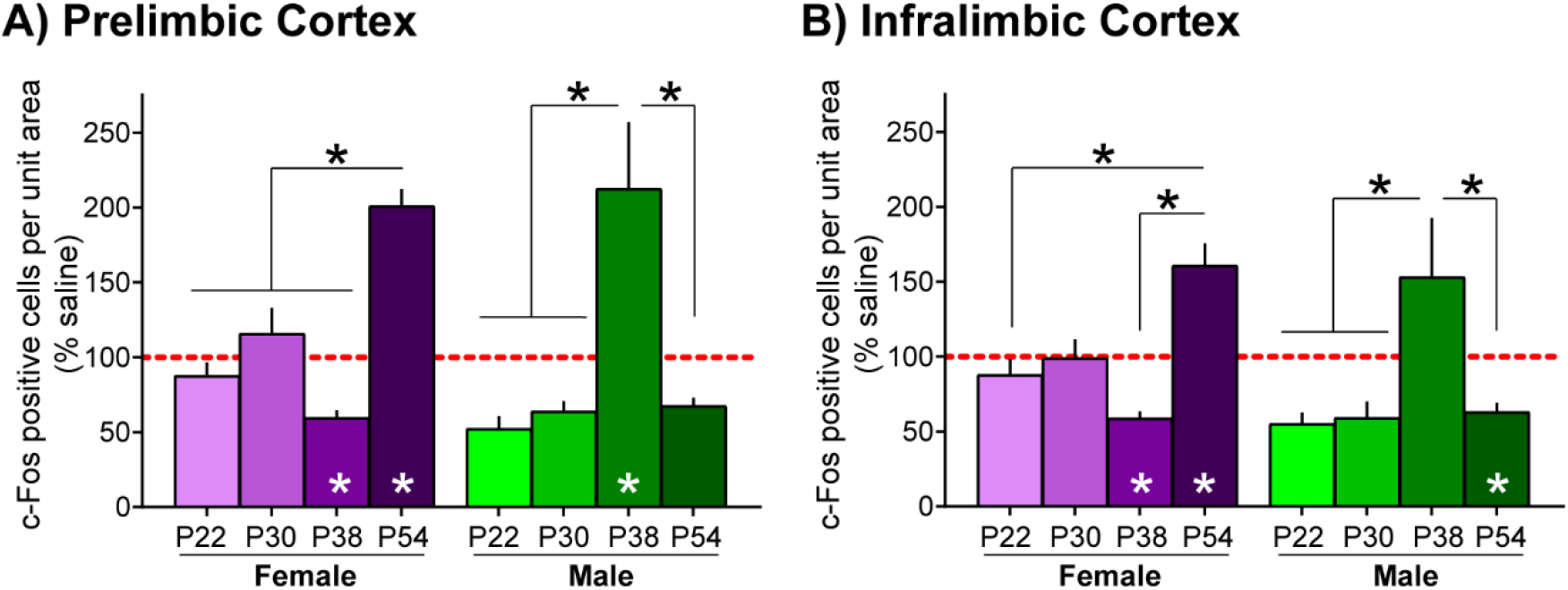
Adolescent age mediates c-Fos expression in PFC after morphine treatment in both sexes. Morphine data were normalized to the average saline response for each age and sex (see Supp. Fig. 1 for saline data). **(A)** In prelimbic cortex, the c-Fos response to morphine treatment at P54 was greater in females than at all other ages. In males, the c-Fos response to morphine at P38 was greater than at all other ages. **(B)** In infralimbic cortex, the c-Fos response to morphine at P54 was greater than P22 and P38 in females. In males, the c-Fos response to morphine at P38 was greater than all other ages. Asterisks at the bottom of each histogram indicate groups in which there was a significant difference between saline and morphine treatments, and asterisks between histograms indicate differences between the c-Fos response to morphine treatment by age (Šídák’s multiple comparisons test; *p*<0.05). *n*=13-16 and *n*=11-13 observations/rat, respectively, *n*=3 rats/group.

**Fig. 2:**
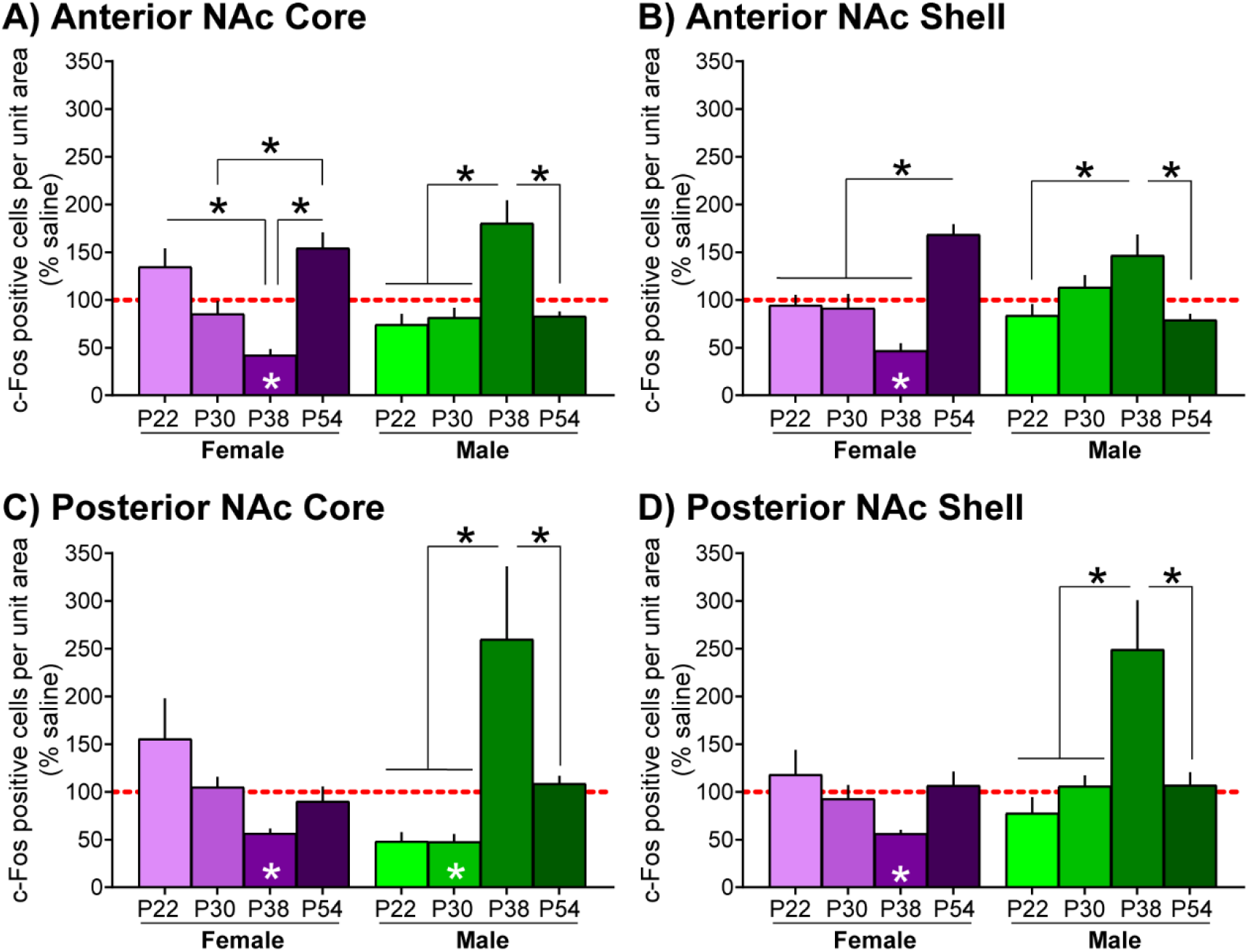
Adolescent age mediates c-Fos expression in NAc after morphine treatment in both sexes. Morphine data were normalized to the average saline response for each age and sex (see Supp. Fig. 2 for saline data). **(A)** In anterior NAc core in females, the c-Fos response to morphine treatment at P38 was less than P22 and P54, and the c-Fos response to morphine at P54 was greater than P30. In males, the c-Fos response to morphine at P38 was greater than at all other ages. **(B)** In anterior NAc shell in females, the c-Fos response to morphine at P54 was greater than all other ages. In males, the c-Fos response to morphine at P38 was greater P22 and P54. **(C)** In posterior NAc core in females, there were no significant differences in the c-Fos response to morphine treatment by age. In males, the c-Fos response to morphine at P38 was greater than at all other ages. **(D)** In posterior NAc shell in females, there were no significant differences in the c-Fos response to morphine treatment by age. In males, the c-Fos response to morphine at P38 was greater than all other ages. Asterisks at the bottom of each histogram indicate groups in which there was a significant difference between saline and morphine treatments, and asterisks between histograms indicate differences between the c-Fos response to morphine treatment by age (Šídák’s multiple comparisons test; *p*<0.05). *n*=15-18, *n*=15-19, *n*=8-13, *n*=10-14 observations/rat, respectively, *n*=3 rats/group.

**Fig. 3:**
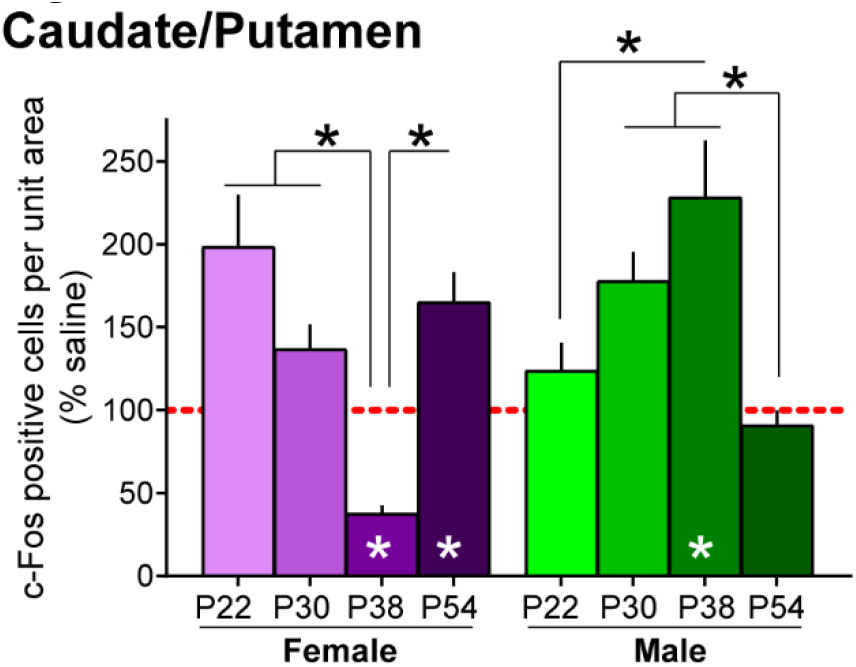
Adolescent age mediates c-Fos expression in CPu after morphine treatment in both sexes. Morphine data were normalized to the average saline response for each age and sex (see Supp. Fig. 3 for saline data). In females, the c-Fos response to morphine treatment at P38 was less than all other ages. In males, the c-Fos response to morphine at P38 was greater than P22, and the c-Fos response to morphine at P54 was less than P30 and P38. Asterisks at the bottom of each histogram indicate groups in which there was a significant difference between saline and morphine treatments, and asterisks between histograms indicate differences between the c-Fos response to morphine treatment by age (Šídák’s multiple comparisons test; *p*<0.05). *n*=23-28 observations/rat, *n*=3 rats/group.

**Fig. 4:**
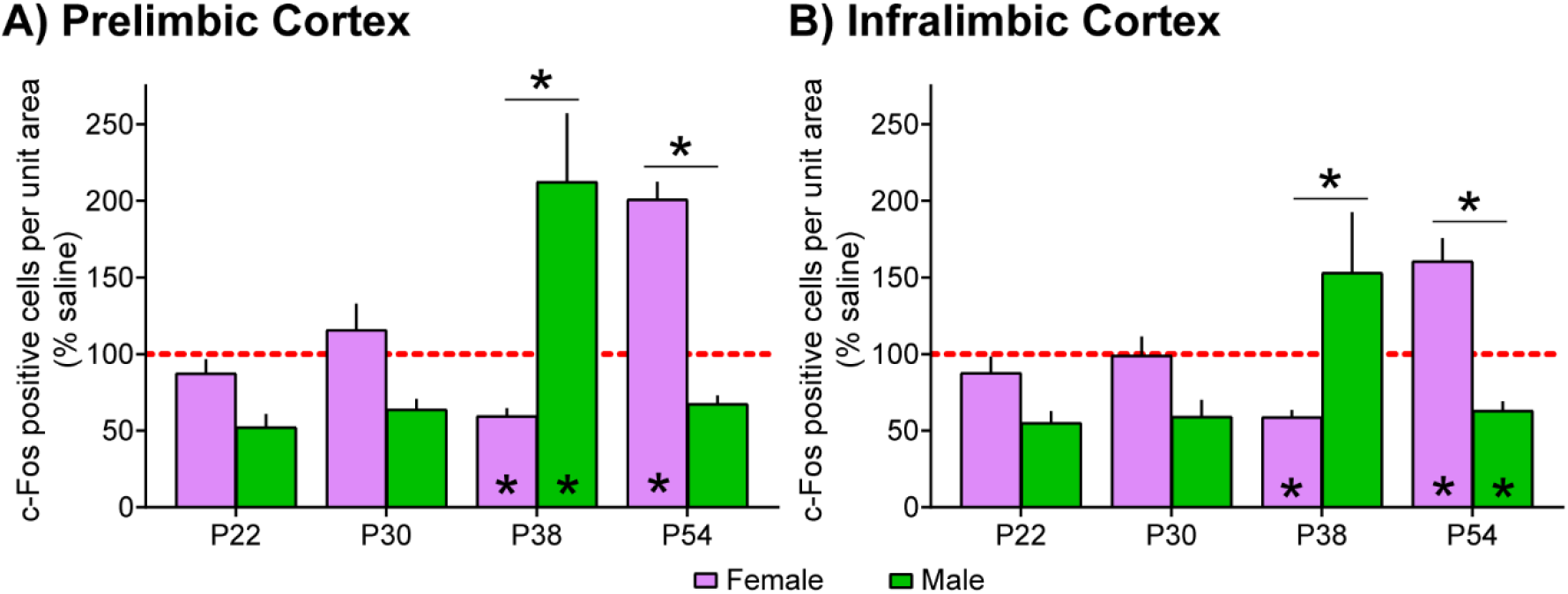
Sex mediates c-Fos expression in PFC after morphine treatment across adolescence. Morphine data were normalized to the average saline response for each age and sex (see Supp. Fig. 1 for saline data). **(A)** In prelimbic cortex, the c-Fos response to morphine treatment was greater in males than in females at P38, and greater in females than males at P54. **(B)** In infralimbic cortex, the c-Fos response to morphine was greater in males than in females at P38, and greater in females than in males at P54. Asterisks at the bottom of each histogram indicate groups in which there was a significant difference between saline and morphine treatments, and asterisks between histograms indicate differences between the c-Fos response to morphine treatment by age (Šídák’s multiple comparisons test; *p*<0.05). *n*=13-16 and *n*=11-13 observations/rat, respectively, *n*=3 rats/group.

**Fig. 5:**
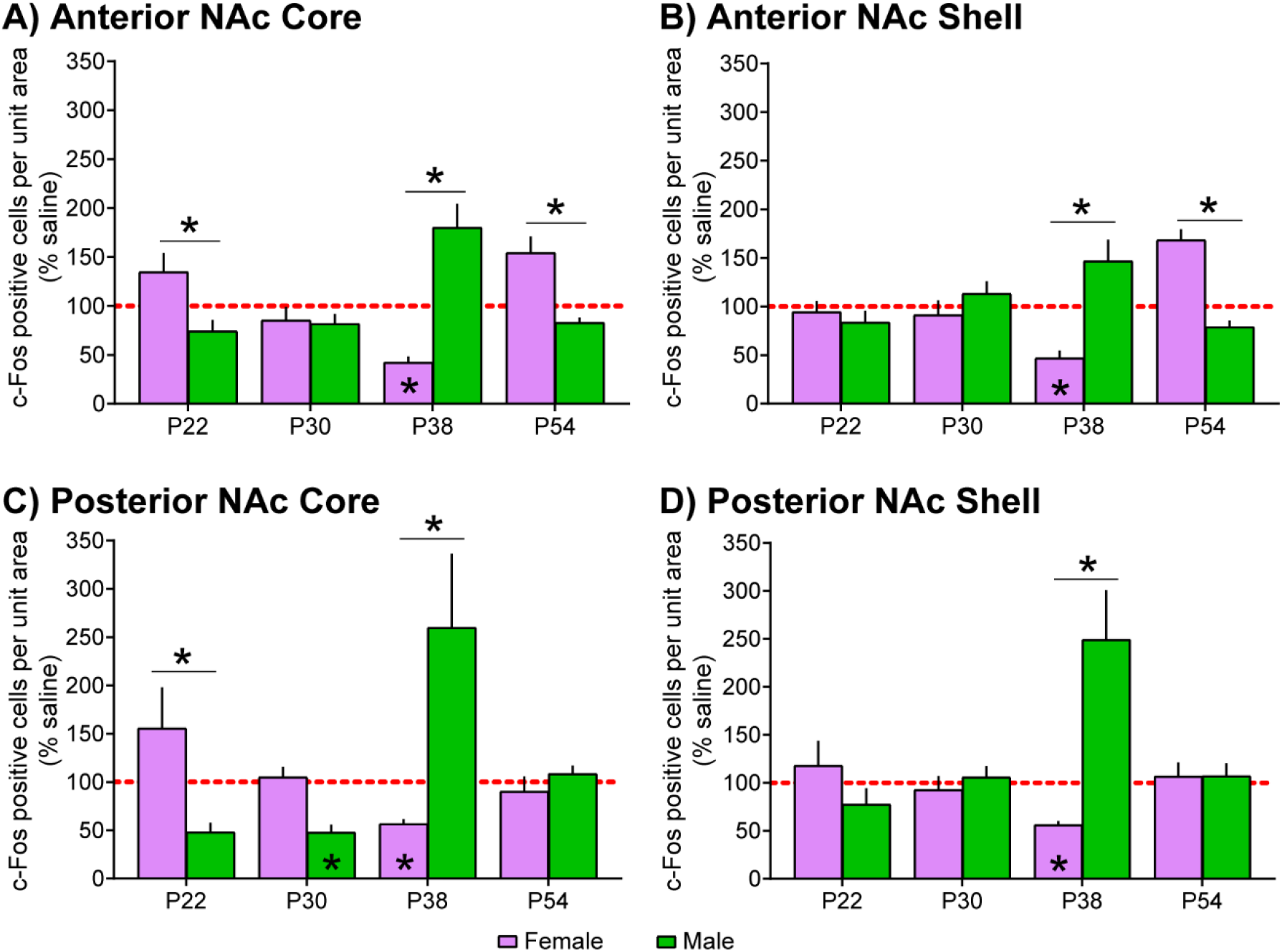
Sex mediates c-Fos expression in NAc after morphine treatment across adolescence. Morphine data were normalized to the average saline response for each age and sex (see Supp. Fig. 2 for saline data). **(A)** In anterior NAc core, the c-Fos response to morphine treatment at P22 and P54 was greater in females than in males, and greater in males than in females at P38. **(B)** In anterior NAc shell, the c-Fos response to morphine was greater in males than in females at P38, and greater in females than in males at P54. **(C)** In posterior NAc core, the c-Fos response to morphine was greater in females than in males at P22, and greater in males than in females at P38. **(D)** In posterior NAc shell, the c-Fos response to morphine was greater in males than in females at P38. Asterisks at the bottom of each histogram indicate groups in which there was a significant difference between saline and morphine treatments, and asterisks between histograms indicate differences between the c-Fos response to morphine treatment by age (Šídák’s multiple comparisons test; *p*<0.05). *n*=15-18, *n*=15-19, *n*=8-13, *n*=10-14 observations/rat, respectively, *n*=3 rats/group.

**Fig. 6:**
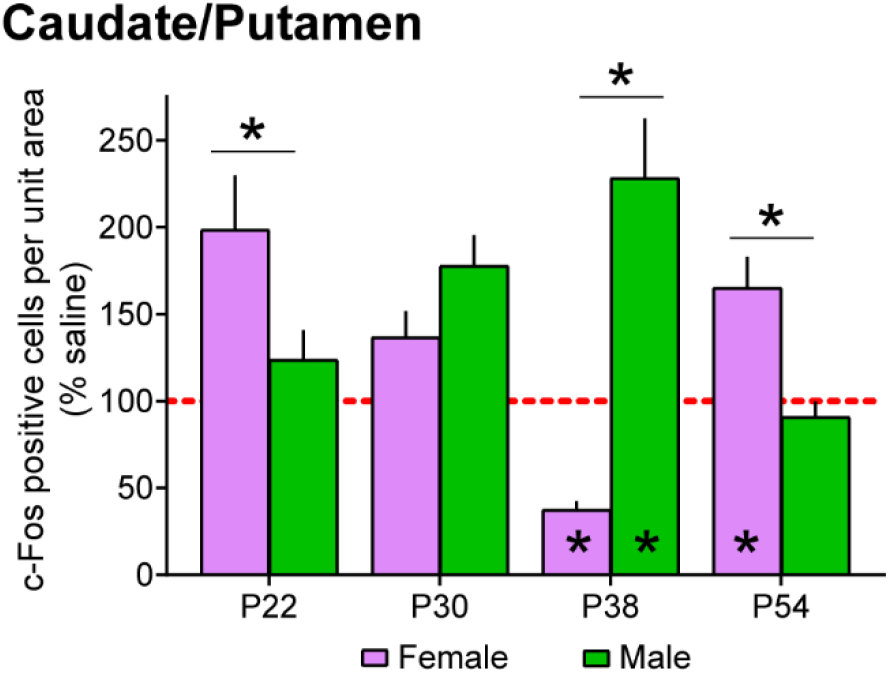
Sex mediates c-Fos expression in CPu after morphine treatment across adolescence. Morphine data were normalized to the average saline response for each age and sex (see Supp. Fig. 3 for saline data). The c-Fos response to morphine treatment was greater in females than in males at P22 and P54, and greater in males than in females at P38. Asterisks at the bottom of each histogram indicate groups in which there was a significant difference between saline and morphine treatments, and asterisks between histograms indicate differences between the c-Fos response to morphine treatment by age (Šídák’s multiple comparisons test; *p*<0.05). *n*=23-28 observations/rat, *n*=3 rats/group.

## Results

We quantified the number of c-Fos positive nuclei in the PFC, NAc, and CPu after morphine (3mg/kg free base) or saline injection in male and female rats. Injections were delivered at either P22, P30, P38, or P54, when we have observed age- and sex-specific behavioral differences and developmental activity in the anterior NAc core [14]. The PFC can be broken into functionally distinct subregions, including the prelimbic cortex and infralimbic cortex [22]. Similarly, the NAc contains functionally distinct core and shell subregions [23]. In both prelimbic and infralimbic cortices and NAc core and shell, there was a significant difference (*p*<0.05) between c-Fos positive cells in each of the subregions (25% of groups; paired two-tailed *t*-tests), and thus we will present data from these regions separately. In NAc, there was also a significant anterior-posterior gradient (25-31% of groups; paired two-tailed *t*-tests), thus we will separate anterior (Bregma +3.24 - +1.8) and posterior (Bregma +1.68 - +0.84) NAc core and shell. An anterior-posterior gradient was present in <25% of groups in prelimbic cortex (0%), infralimbic cortex (6%), and CPu (19%), thus we grouped all data for these regions.

There were changes in the number of c-Fos positive cells in saline-injected rats across adolescence (**Supp. Table 1**), which could be due either to basal changes in neural activity in the regions examined or to changes in sensitivity to injection stress. Because all ROIs examined undergo developmental plasticity during adolescence, it is plausible that there are baseline changes in spontaneous neural activity across different adolescent stages [24, 25]. Without a naïve control group we cannot say for certain, and thus for clarity of interpretation we depicted morphine data as a percentage of the average saline response, with 100% indicating no change from saline-treated rats. There was a significant interaction between drug, sex, and age in all regions (all *p*<0.001; prelimbic: *F*_(3, 217)_=16.59, infralimbic: *F*_(3,190)_=9.23, anterior NAc core: *F*_(3, 231)_=11.15, anterior NAc shell: *F*_(3,208)_=6.21, posterior NAc core: *F*_(*3,144*)_=*10.39*, posterior NAc shell: *F*_(3,165)_=6.42, CPu: *F*_(3, 355)_=22.18), and thus we will plot the same data in two different ways to highlight age-dependent differences or sex-dependent differences.

### The effect of morphine on c-Fos expression

Significant differences between saline and morphine groups are indicated in Figs. 1-6 as asterisks within individual histograms. Full data sets with both saline and morphine histograms can be found in **Supp. Figs. 1-3**. Within the prelimbic cortex in females, morphine treatment significantly reduced c-Fos expression relative to saline-injected controls at P38 (*p*=0.002), but increased c-Fos at P54 (*p*<0.001). In male prelimbic cortex, the only statistically significant effect of morphine treatment was increased c-Fos at P38 (*p*=0.003; **Fig. 1A, 4A, Supp. Fig. 1**). Within the infralimbic cortex in females, morphine reduced c-Fos at P38 and increased c-Fos at P54 (*p*=0.005, 0.014, respectively), similar to prelimbic cortex patterns. Within the infralimbic cortex in males, morphine significantly reduced c-Fos at P54 (*p*=0.048; **Fig. 1B, 4B, Supp. Fig. 2**). Within the anterior NAc, the only significant effects of morphine treatment on c-fos expression were significant reductions in both the anterior NAc core and shell in females at P38 (*p*<0.001, 0.003, respectively; **Fig. 2A-B, 5A-B, Supp. Fig. 2A-B**). Within the posterior NAc, morphine treatment significantly reduced c-Fos at P38 in females in both the core and shell (both *p*<0.001). In males, morphine treatment significantly reduced c-Fos at P30 in the posterior NAc core (*p*=0.018), but not the shell (**Fig. 2C-D, 5C-D, Supp. Fig. 2C-D**). Within the CPu, morphine treatment significantly reduced c-Fos expression at P38, and significantly increased c-Fos at P54 in females (*p*<0.001, =0.002, respectively). In male CPu, morphine significantly increased c-Fos at P38 (*p*=0.007; **Fig. 3, 6, Supp. Fig. 3**).

### The effect of adolescent age on the c-Fos response to morphine

Significant differences between adolescent ages are depicted as asterisks between histograms in Figs. 1-3. Within the prelimbic cortex in females, c-Fos expression in morphine-treated relative to saline-treated rats was significantly greater at P54 than all other adolescent ages (P22: *p*<0.001, P30: *p*=0.020, P38: *p*<0.001), while in males c-Fos expression in response to morphine was significantly greater at P38 than all other ages (all *p*<0.001; **Fig. 1A**). Within the infralimbic cortex, c-Fos expression in response to morphine at P54 was significantly greater than at P22 and P38 (P22: *p*=0.030, P38: *p*<0.001), while in males the morphine response at P38 was greater than all other ages (P22: *p*<0.001, P30: *p*=0.002, P54: *p*<0.002; **Fig. 1B**). Within the anterior NAc core in females, c-Fos expression in response to morphine was significantly lower at P38 than at P22 or P54 (both *p*<0.001), and P54 responses were also higher than P30 (*p*=0.010). Within the anterior NAc core in males, c-Fos expression in response to morphine was significantly greater at P38 than all other adolescent ages (all *p*<0.001; **Fig. 2A**). Within the anterior NAc shell in females, c-Fos expression in response to morphine was significantly greater at P54 than all other ages (P22: *p*=0.003, P30: *p*<0.001, P38: *p*<0.001), while in males the response at P38 was significantly greater than at P22 or P54 (P22: *p*=0.005, P54: *p*=0.002; **Fig. 2B**). There were no significant changes in c-Fos expression in response to morphine across adolescent age in females within the posterior NAc core or shell in females, but in males the response at P38 was greater than all other ages in both the posterior core (P22: *p*<0.001, P30: *p*<0.001, P54: *p*=0.001) and the shell (all *p*<0.001; **Fig. 2C-D**). Within the female CPu, c-Fos expression at P38 was significantly decreased relatively to all other ages assessed (P22: *p*<0.001, P30: *p*=0.003, P54: *p*<0.001). In the male CPu, P38 c-Fos induction was significantly increased relative to P22 (*p*=0.002), while both P30 and P38 c-Fos induction was significantly increased relative to P54 (P30: *p*=0.009, P38: *p*<0.001; **Fig. 3**).

### The effect of sex on the c-Fos response to morphine

Significant differences between sexes are depicted as asterisks between histograms in Figs. 4-6. Within both the prelimbic and infralimbic cortices, there was a significantly different c-Fos response to morphine between males and females at P38 (males>females), which reversed at P54 (females>males) (all *p*<0.001; **Fig. 4**). Within the anterior NAc core, females had a greater c-Fos response to morphine than males at P22 (*p*=0.021) and P54 (*p*=0.005), but males had a greater response at P38 (*p*<0.001; **Fig. 5A**). In the anterior NAc shell, males had a greater c-Fos response at P38, but females had a greater response at P54 (both *p*<0.001; **Fig. 5B**). In posterior NAc core, females had a greater c-Fos response to morphine at P22 (*p*=0.044), and males had a greater response at P38 (*p*<0.001; **Fig. 5C**). In posterior NAc shell, males had a greater response at P38 (*p*<0.001; **Fig. 5D**). And within the CPu, females had a greater c-Fos response to morphine than males at P22 (*p*=0.042) and P54 (*p*=0.036), and males had a greater response than females at P38 (*p*<0.001; **Fig. 6**).

## Discussion

Adolescence is a known critical period of development for the PFC, NAc, and CPu. Our data add to this body of literature that there may be stages *within* adolescence that are more responsive to opioids and, moreover, those sensitive periods may be sex-specific. There are data from rats that indicate males are more sensitive to a mid-adolescent ethanol exposure than females, and that this sensitivity is absent at late adolescence [13, 26]. Similarly, repeated morphine exposure during mid-adolescence, but not late adolescence, results in dysfunctional reward-related behavior later in life (females not assessed) [27]. Age during adolescence and sex are also factors in drug taking behavior in humans [12, 28], and in the likelihood that opioid prescriptions are filled post-surgery during adolescence [6].

### Direction of c-Fos regulation by morphine (Fig. 7)

Morphine is often reported to increase c-Fos expression [19, 29, 30], and only in the context of nociception (i.e. morphine analgesia) is a decrease in c-Fos expression commonly reported [31, 32]. Only one other study to our knowledge has reported decreased c-Fos induction after morphine administration with no pain-related stimuli, which was dependent on the novelty of the environment in which morphine was administered [33]. Our data expand this notion by demonstrating that morphine dynamically alters c-Fos induction in different developmental, sex, and neuroanatomical contexts. Morphine produced both increased and decreased c-Fos expression in the regions we assessed. Other addictive substances have also been reported to reduce c-Fos expression in some brain regions, while increasing c-Fos in others [34, 35]. Regardless of direction, significant effects of morphine (relative to saline) on c-Fos expression were more pronounced in females than in males. In females, acute morphine exposure at P38 significantly reduced c-Fos in all regions assessed. In posterior NAc and CPu, this may be a function of increased c-Fos in saline-injected rats relative to other ages (**Supp. Table 1**), which was then suppressed by morphine (**Supp. Fig. 2, 3**). However, in PFC and anterior NAc, c-Fos in saline-injected rats was not significantly different from all other ages (**Supp Table 1**; **Supp. Fig. 1, 2**). Males also exhibited differences in c-Fos induction across adolescence in saline-treated groups (**Supp. Table 1**). Spontaneous neural activity is common in developing brain regions [24, 25]. Thus, we predict that there is a basal increase in neural activity in some brain regions at P38 in females rather than a sensitized stress response to injection, though we cannot rule out the latter at this time. In female PFC and CPu, the suppressive effects of morphine at P38 were reversed at P54, at which time morphine significantly increased c-Fos. Female puberty occurs ~P35 in rats [36], and both estradiol and estrous stage have been demonstrated to modify the analgesic responses to morphine in rats [37, 38]. Opioid treatment can also alter endocrinological parameters in both male and female rats [39–41]. However, not all sex differences in reward circuitry function can be explained by puberty [42, 43], and there may be species-specific sex differences [44]. Future studies should examine how the induction of cycling sex hormones impacts dynamic changes in morphine-induced neural activity.

**Fig. 7:**
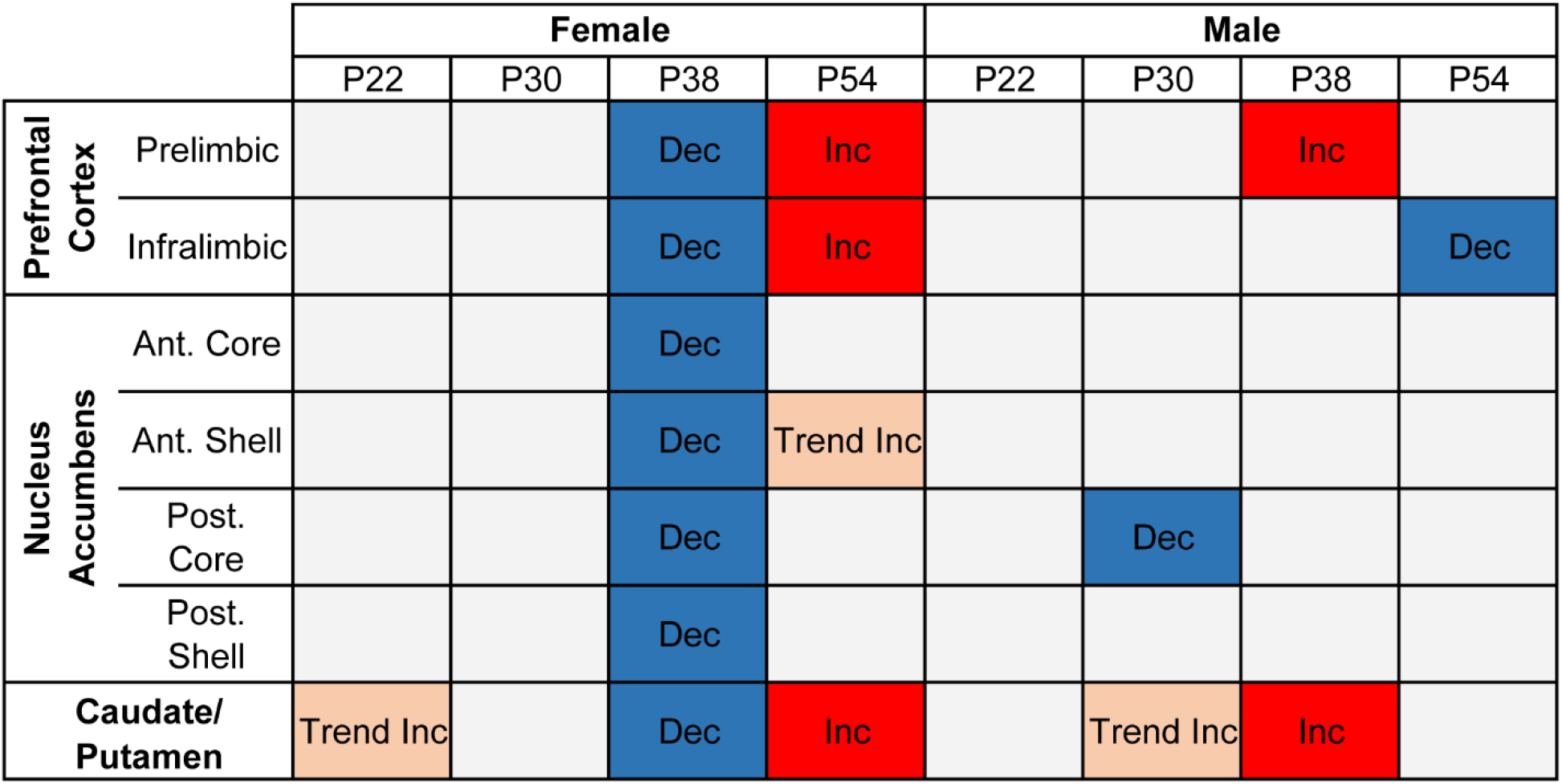
Summary of the statistically significant effects of morphine (relative to saline) on c-Fos expression across adolescence in each sex. Significant (Šídák’s multiple comparisons test; *p*<0.05) decreases in c-Fos expression in response to morphine vs. saline treatment are indicated with blue squares and “Dec”. Significant (Šídák’s multiple comparisons test; *p*<0.05) increases in c-Fos expression in response to morphine vs. saline treatment are indicated with red squares and “Inc”. Trends toward increased c-Fos expression in response to morphine vs. saline (Šídák’s multiple comparisons test; *p*<0.10) are indicated in pink and “Trend Inc”.

### Region-specific effects of morphine across adolescence (Fig. 8)

Our data suggest there are sex- and region-specific patterns of neural responding to morphine across adolescence. In males, there was a transient increase in morphine-induced c-Fos expression in PFC and NAc at P38 relative to all other ages. Male CPu was distinct in that morphine-induced c-Fos expression elevation was prolonged, and increased at both P30 and P38 relative to other ages. Female effects were more varied. Morphine-induced c-Fos expression in female PFC and anterior NAc shell was higher at P54 than most other ages, and there were no significant effects in posterior NAc. In female NAc core and CPu there was a transient decrease in morphine-induced c-Fos expression at P38 relative to most other ages. Taken together, these data suggest that developmental transitions between P30-38 and P38-54 may be particularly sensitive to the effects of opioid exposure in *both* sexes, but sex differences in the direction of neural responding to morphine (discussed further in the next section), suggest that opioid exposure will produce opposing behavioral effects between the sexes. Finally, each region examined was selected because it undergoes documented developmental plasticity during adolescence, notably synaptic pruning [14, 45–50]. Each region also contributes to reward-related behavior in a unique way. Broadly speaking, the prelimbic cortex is associated with reward seeking, while the infralimbic cortex is associated with reward-related behavioral extinction [22]. Our data indicate that the prelimbic and infralimbic cortices largely respond to morphine across adolescence in the same, sex-specific patterns. The NAc core is associated with reward-mediated approach and learning, while the NAc shell is associated with activating patterns of behavior that result in reward, and suppressing interfering behavioral patterns [23]. Our data indicate that morphine alters NAc neural activity differently in anterior vs. posterior NAc, an effect that is more prominent in females than in males. An anterior-posterior NAc gradient is well-documented in the literature [51–55], including a sex-specific gradient [56], and there are fMRI studies in humans that support a functional dissociation between anterior and posterior striatal regions [57–59]. Lastly, the CPu is important for habit formation and motor responses [60]. Our data suggest that there may be some fundamental differences in the developmental trajectory of the CPu relative to other reward-related regions in both sexes. Adolescent drug exposures can increase substance use disorders later in life [2, 3, 5–7], which have symptoms related to both habit and reward [61, 62]. These data may suggest that adolescent drug exposure could influence habit- and reward-dysfunction via independent, but parallel developmental abnormalities.

**Fig. 8:**
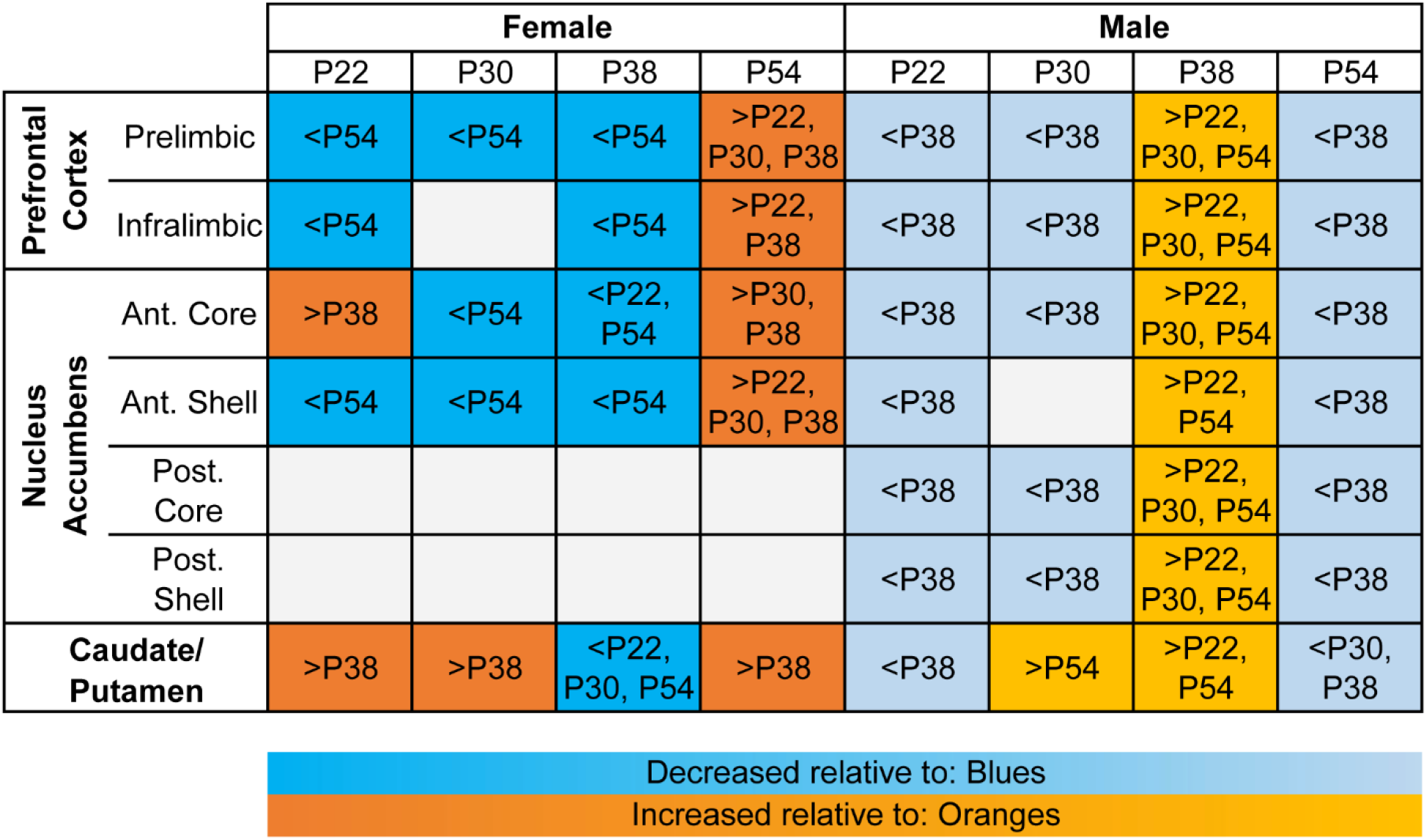
Summary of the statistically significant effects of adolescent age on morphine-induced c-Fos expression in each sex. Significant (Šídák’s multiple comparisons test; *p*<0.05) decreases in c-Fos expression in response to morphine vs. saline treatment are indicated with blue squares, and the text indicates to which age the response is different. Significant (Šídák’s multiple comparisons test; *p*<0.05) increases in c-Fos expression in response to morphine vs. saline treatment are indicated with orange squares, and the text indicates to which age the response is different.

### Sex differences in the neural response to morphine (Fig. 9)

Sex differences have been observed in adolescent development [12, 14] and in response to opioids in both pain/nociception- [63, 64] and substance use disorder-related contexts [15, 28]. Nearly all ROIs examined had a sex difference in response to morphine in both directions, male>female and female>male, over the course of adolescence. In all ROIs, males had a greater c-Fos response to morphine than females at P38. In all regions except posterior NAc, this reversed at P54, when females had a greater c-Fos response than males. At P22, females had a greater c-Fos response than males in the NAc core and CPu. These data indicate that some sex differences exist prior to puberty, and thus are unlikely to be due to cycling sex hormones. There are data indicating that early postnatal sex hormone signaling can regulate adolescent reward-related neural function and behavior in sex-specific ways [16, 42, 65], thus it remains possible these sex differences occurred due to androgen programming around birth. We have also observed that pruning-related activity in the NAc core targets sex-specific synaptic proteins and occurs prior to puberty in both sexes, but earlier in females than in males (P22 vs. P30, respectively) [14]. It is unclear how sex-specific NAc pruning might alter the neural activity response to morphine.

**Fig. 9:**
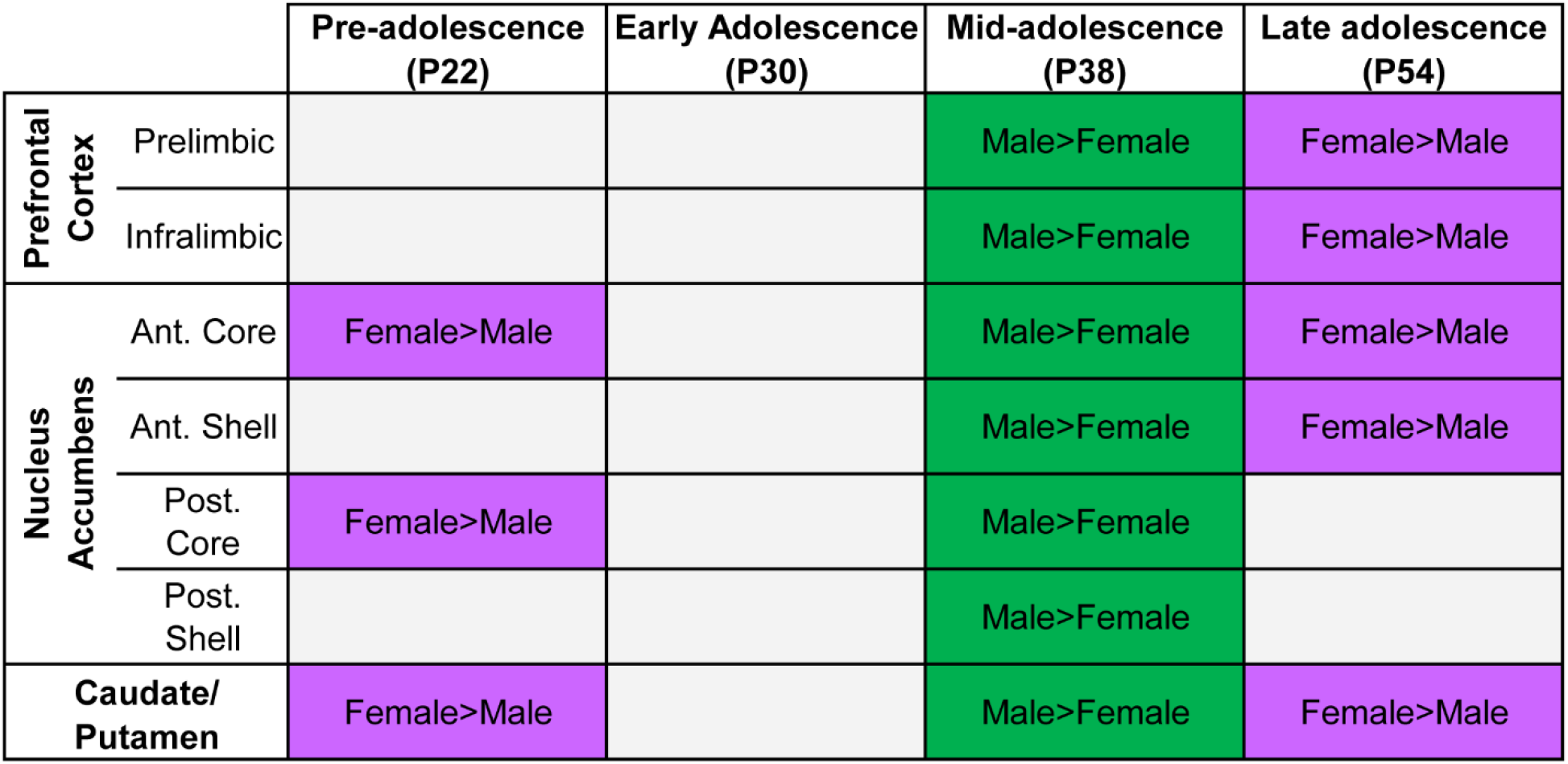
Summary of the statistically significant effects of sex on morphine-induced c-Fos expression across adolescence. Significant (Šídák’s multiple comparisons test; *p*<0.05) increases in female c-Fos expression in response to morphine vs. saline treatment relative to males are indicated with purple squares and “Female>Male”. Significant (Šídák’s multiple comparisons test; *p*<0.05) increases in male c-Fos expression in response to morphine vs. saline treatment relative to females are indicated with green squares and “Male>Female”.

In summary, we propose it will be important in future studies to add age within adolescence as an independent variable to fully capture the consequences of healthy or abnormal reward-related neural development.

## Acknowledgments

We thank the Pathology Clinical Research Core at Albany Medical Center for assistance with imaging. This work was supported by F32DA043308, R03AG070111, and Albany Medical College start-up funds to AMK and R01DA034185 and R01MH101183 to SDB.

---

**Supp. Table 1:** Statistically significant differences in c-Fos expression across adolescence in vehicle-treated rats. Šídák’s multiple comparisons tests; ANOVA results and group sizes for each region indicated in Supp. Figs. 1-3. No text indicates the comparison was non-significant (*p*>0.05).

|  |  | Prelimbic | Infralimbic | Ant. NAc<br>core | Ant. NAc<br>shell | Post. NAc<br>core | Post. NAc<br>shell | Cpu |
| --- | --- | --- | --- | --- | --- | --- | --- | --- |
| Female | P22:P30 | | $p=0.030$ | $p=0.001$ | | | | |
| | P22:P38 | $p<0.001$ | $p=0.025$ | $p<0.001$ | $p=0.002$ | $p<0.001$ | $p<0.001$ | $p<0.001$ |
| | P22:P54 | | | | | | | $p=0.045$ |
| | P30:P38 | | | | | $p<0.001$ | $p=0.005$ | $p<0.001$ |
| | P30:P54 | | $p=0.020$ | | | | | |
| | P38:P54 | $p<0.001$ | $p=0.016$ | $p<0.001$ | $p=0.005$ | $p<0.001$ | $p<0.001$ | $p<0.001$ |
| Male | P22:P30 |  |  |  |  |  |  |  |
| | P22:P38 | | | | | | $p=0.019$ | |
| | P22:P54 | $p<0.001$ | $p=0.027$ | $p=0.031$ | | | | $p<0.001$ |
| | P30:P38 | | | | | $p<0.001$ | | |
| | P30:P54 | | | $p=0.015$ | $p=0.023$ | | | $p<0.001$ |
| | P38:P54 | $p<0.001$ | $p<0.001$ | $p<0.001$ | $p<0.001$ | | | $p<0.001$ |

**Supp. Fig. 1:**
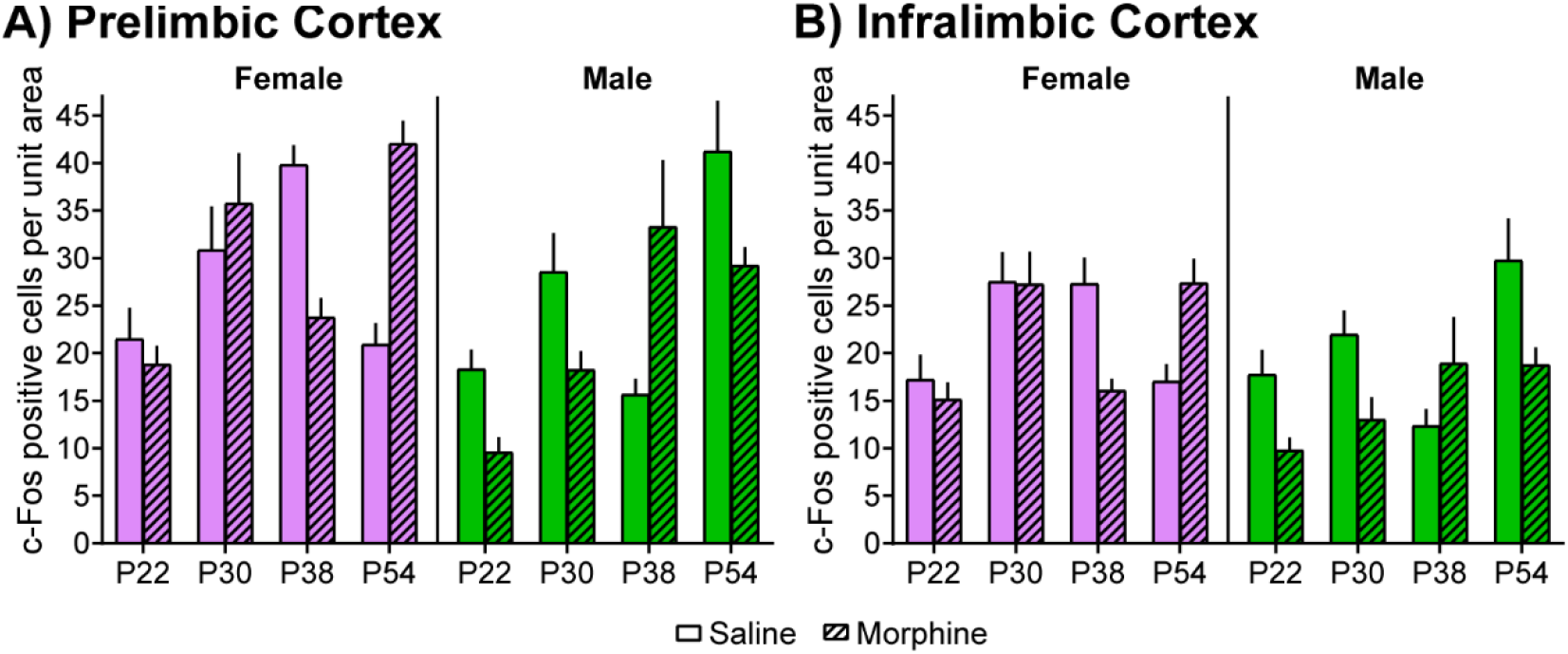
c-Fos expression in PFC after saline or morphine treatment across adolescence in both sexes. There was a significant interaction (*p*<0.05) between age, sex, and drug in both **(A)** prelimbic cortex (*F*_(3, 217)_=16.59) and **(B)** infralimbic cortex (*F*_(3,190)_=9.23). Post-hoc test results are indicated in the main figures (Fig. 1,4). *n*=13-16 and *n*=11-13 observations/rat, respectively, *n*=3 rats/group.

**Supp. Fig. 2:**
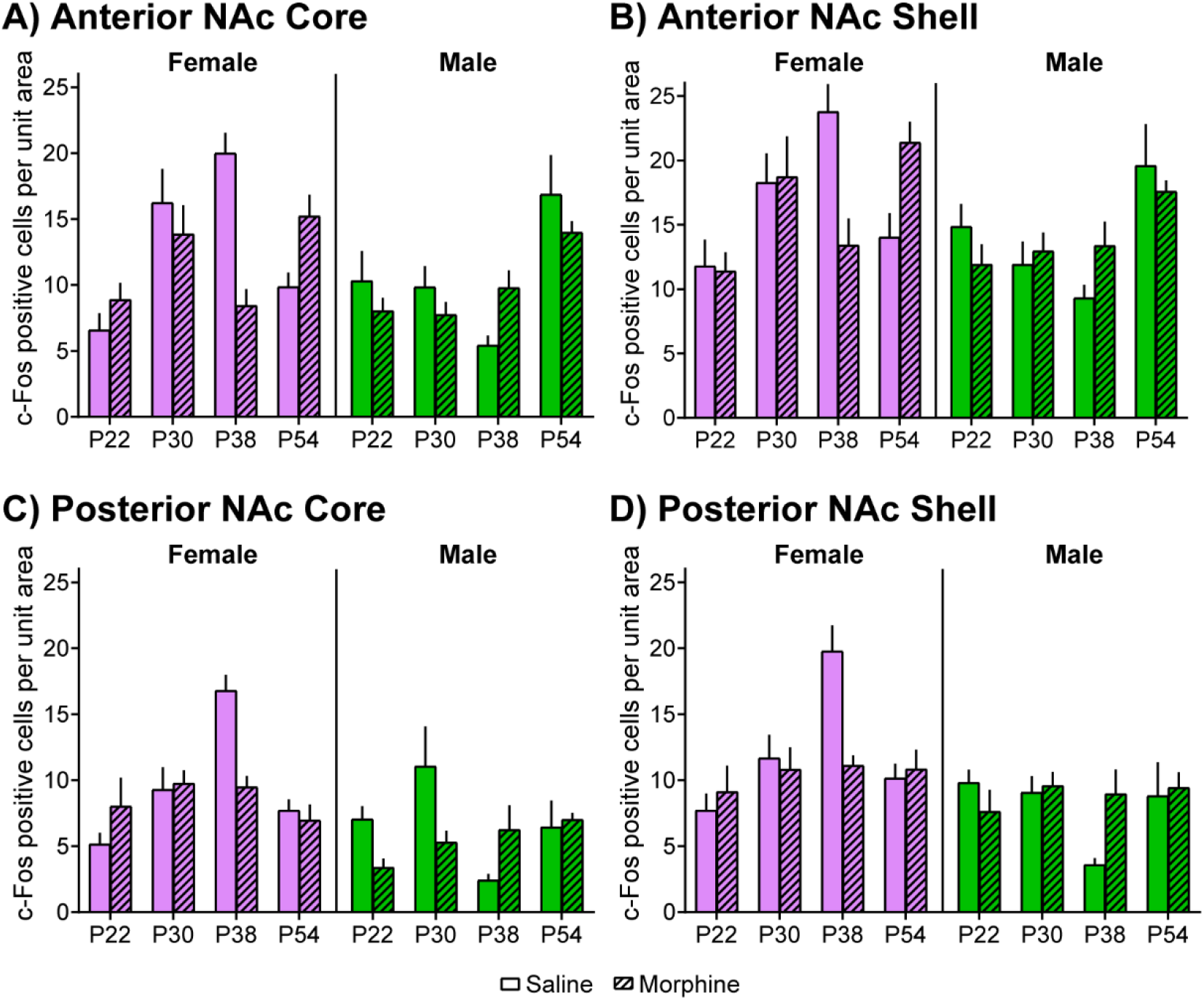
c-Fos expression in NAc after saline or morphine treatment across adolescence in both sexes. There was a significant interaction (*p*<0.05) between age, sex, and drug in **(A)** anterior NAc core (*F*_(3, 231)_=11.15), **(B)** anterior NAc shell (*F*_(3,208)_=6.21), **(C)** posterior NAc core (*F*_(3,144)_=10.39), and **(D)** posterior NAc shell (*F*_(3,165)_=6.42). Post-hoc test results are indicated in the main figures (Fig. 2,5). *n*=15-18, *n*=15-19, *n*=8-13, *n*=10-14 observations/rat, respectively, *n*=3 rats/group.

**Supp. Fig. 3:**
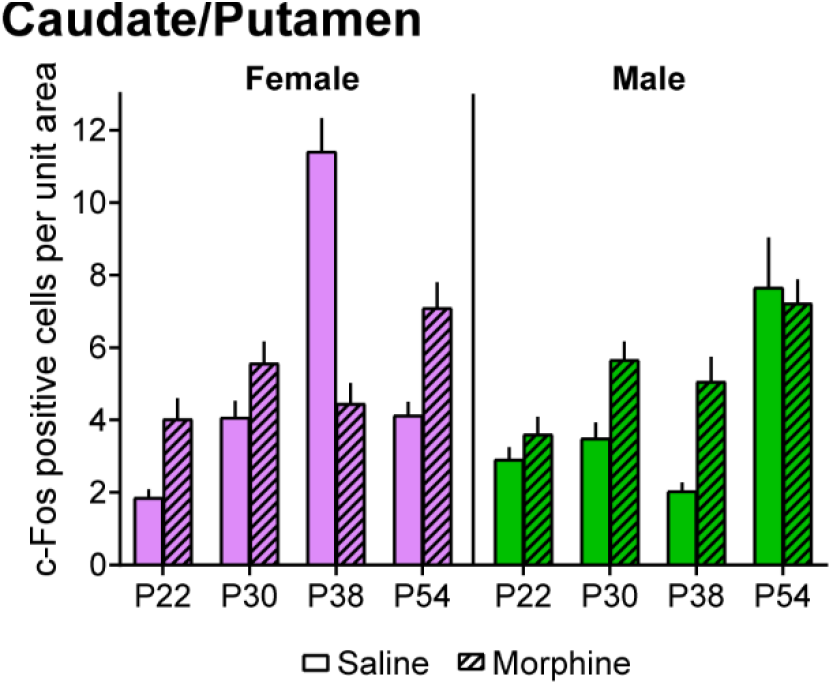
c-Fos expression in CPu after saline or morphine treatment across adolescence in both sexes. There was a significant interaction (*p*<0.05) between age, sex, and drug in CPu (*F*<_(3, 355)_=22.18). Post-hoc test results are indicated in the main figures (Fig. 3,6). *n*=23-28 observations/rat, *n=3* rats/group.

## Notes

### Competing Interest Statement

The authors have declared no competing interest.

